# A simple centrifugation protocol leads to a 55-fold mitochondrial DNA enrichment and paves the way for future mitogenomic research

**DOI:** 10.1101/106583

**Authors:** Jan Niklas Macher, Vera Zizka, Alexander Martin Weigand, Florian Leese

**Author notes:** JNM, >.

## Abstract

DNA (meta)barcoding is used to study biodiversity and is available for standardised assessments. However, it suffers from PCR bias, which can lead to the loss of specific taxa. PCR-free techniques such as shotgun metagenomics are therefore thought to be more suited for biodiversity assessments, but are currently limited by incomplete reference libraries.

The technique of ‘mitogenome-skimming’ or ‘mitogenomics’, in which complete mitochondrial genomes are sequenced, is ideal to bridge the techniques of (meta)barcoding and metagenomics. However, without the enrichment of mitochondria, roughly 99 % of all sequencing reads are of non-mitochondrial origin and mostly useless for common applications, e.g. species identification.

Here, we present a simple centrifugation protocol that leads to an average 140-fold enrichment of mitochondrial DNA. By sequencing six ‘mock’- communities – comprising the freshwater taxa *Corbicula fluminea, Gammarus roeselii* and *Hydropsyche exocellata* each – we recovered whole mitochondrial genomes of these species and the acanthocephalan endoparasite *Pomphorhynchus laevis*.

The presented protocol will greatly speed up building reference libraries for whole mitochondrial genomes, as dozens of species could be sequenced on a single MiSeq run. Subsequently, it will also allow biodiversity assessments using mitogenomics at greatly reduced costs in comparison to mitogenomic approaches without prior enrichment for mitochondria.

## Introduction

Biodiversity is highly important for intact ecosystems and inevitable for human well being (Rockström *et al*. 2009). Molecular techniques such as DNA barcoding (Hebert *et al*. 2003) and metabarcoding (Hajibabaei *et al*. 2011) are increasingly used for biodiversity research but suffer from PCR stochasticity and primer bias (Elbrecht and Leese 2015). The same bias can be introduced by the use of baits or probes (e.g. (Liu *et al*. 2015); (Mayer *et al*. 2016). Therefore, PCR and primer/probe-free techniques harbor the potential for future biodiversity assessments (Tang *et al*. 2014; Zhou *et al*. 2013; Elbrecht and Leese 2015; Crampton-Platt *et al*. 2016; Coissac *et al*. 2016), by circumventing taxon-dependent PCR amplification biases and offering the possibility to correlate read numbers with biomasses. Since reference libraries are still largely incomplete for nuclear genomic information, but relatively comprehensive for mitochondrial genes, such as the cytochrome c oxidase subunit I (COI) gene for animals, the consequent step towards a PCR-free analysis of biodiversity samples could be seen in ‘mitochondrial metagenomics’, ‘mitogenomics’ or ‘mitogenome-skimming’ (e.g. Tang et al. 2014; Crampton-Platt *et al*. 2015). This technique enables the comparison of newly generated mitogenomes or mitogenome fragments with reference databases and thereby links genomic information to taxonomic knowledge. However, the currently applied approaches are relatively ineffective in terms of sequencing capacity, with most ‘PCR-free’ mitogenomic libraries comprising less than 1 % of sequence reads of mitochondrial origin (Crampton-Platt *et al*. 2016). The major methodological disadvantage thus is the great sequencing depth needed, and the associated high costs for those approaches.

Enrichment of mitochondria prior to DNA extraction and library sequencing is a potential solution, shifting the initial ratio of mitochondrial versus nuclear DNA towards a higher mitochondrial DNA proportion. It is known that ultracentrifugation in CsCl-gradients can enrich for the typically AT-rich mitochondrial genomes (e.g. Garber and Yoder 1983). However, this approach grounds on cost- and labor-intensive ultracentrifugation and mitochondria can have highly variable AT-contents, rendering extractions from bulk biodiversity samples in CsCl-gradients less straight-forward. The enrichment of mitochondria is most promising when organelles are intact, i.e. when living tissue is used (Tamura and Aotsuka 1988). Until now, this approach has not been tested for environmental bulk samples, mainly because most specimens used for biodiversity assessments and genome sequencing are commonly stored in preservation fluids, which damage or destroy mitochondria. Here, we use a ‘mock’- community of three freshwater species to test a simple centrifugation protocol for mitochondrial enrichment in a metagenomic context. We demonstrate that our protocol strongly enriches mitochondrial DNA and therefore can greatly reduce costs of future mitogenomic approaches, e.g. when a) constructing mitochondrial reference libraries and b) assessing biodiversity by an approach which omits biases introduced by primers, probes and PCR reactions.

## Material and Methods

### Sampling and laboratory protocols

Sampling was conducted at two locations (51°00′52.6″N 6°41′04.5″E; 51°05′23.4″N 6°41′17.0″E) of the Gillbach (Germany) in December 2016. Twenty individuals of each of the three macrozoobenthic freshwater species *Corbicula fluminea, Gammarus roeselii* and *Hydropsyche exocellata* were sampled with a dip net or collected from stones. Specimens were transferred into water (500 mL) and transported to the laboratory for immediate processing. Specimens were weighed (Mettler Toledo XS105, table S1) and assembled to six ‘mock’-communities, each containing three individuals of *G. roeselii* and *H. exocellata* and a single *C. fluminea* specimen. ‘Mock’-communities were separately transferred into 3 mL 5° C cold homogenization buffer (0.25 M sucrose, 10 mM EDTA, 30 mM Tris-HCl, pH 7.5; (Tamura and Aotsuka 1988) in a mortar and crushed with a pestle until tissue was homogenized (70 strokes each). Two millilitres of homogenate was pipetted into a 2 mL Eppendorf tube and samples were treated after the following centrifugation protocols (4° C, centrifuge Eppendorf 5427 R) (see Supplementary material 1 for short protocol).

1. Samples 1-3 (‘Complete’- no enrichment of mitochondria): samples were centrifuged for 1 minute at 1,000 g. This step was repeated four times. Final centrifugation was conducted for 10 minutes at 14,000 g. Supernatant was discarded and 600 μL TNES (50 mM Tris Base, 400 mM NaCl, 20 mM EDTA, 0.5 % SDS) buffer was added to the pelleted material. Samples were then homogenized by vortexing.
2. Samples 4-6 (‘Mito’- enrichment of mitochondria): samples were centrifuged for 1 minute at 1,000 g. Pelletized material was discarded, the supernatant transferred to a new tube and again centrifuged for 1 minute at 1,000 g. This step was repeated three times. Final centrifugation was conducted for 10 minutes at 14,000 g. Supernatant was discarded, 600 μL TNES buffer was added to the pelleted material and samples were homogenized by vortexing.

A total volume of 40 μL Proteinase K (300 U/ml, 7Bioscience, Hartheim, Germany) was added to each sample, which were than vortexed and incubated at 37° C for 12 hours (Eppendorf Thermomixer C). DNA was extracted using a salt precipitation protocol as in (Weiss and Leese 2016). For RNA digestion, 1.5 μL RNAse (1.5 μg, Thermo Fisher Scientific, Oberhausen, Germany) was added to each reaction and incubated at 34° C for 30 min on a Thermomixer, followed by a clean up using the MinElute Reaction CleanUp Kit (Qiagen, Hilden, Germany). For DNA fragmentation, samples were placed in an ultrasonic bath (Bandelin SONOREX, RK 510 Hz) for 8 hours. Library preparation was performed with a TruSeq Nano DNA LT Library Prep Kit (Set A, step (2) ‘Repair Ends and Select Library Size’ – (5) ‘Enrich DNA Fragments’). After each step, fragment lengths and concentrations were quantified on a Fragment Analyzer (Advanced Analytical, Automated CE Systems). Samples were equimolar pooled and sent for sequencing on a MiSeq sequencer (v2 chemistry, 250 bp paired-end) at GATC-Biotech (Konstanz, Germany).

### Sequence analysis

Raw sequences were checked for remaining adapters and trimmed with BBDuk as implemented in Geneious v.10.0.9 (Kearse *et al*. 2012). The complete mitochondrial genomes of *Corbicula fluminea, Gammarus roeselii, Hydropsyche exocellata* and *Pomphorhynchus laevis* (an acanthocephalan endoparasite) were assembled using MIRA 4.0.2 (Chevreux, 2014) as implemented in Geneious (Settings: Genome, accurate). Contigs were elongated by mapping reads against contigs with the Geneious mapper (Settings: no gaps allowed, 3% maximum mismatch per read, word length 40). Annotations were performed with the MITOS (Bernt *et al*. 2013) web server and corrected manually.

Analyses of mitochondrial enrichment were conducted after quality filtering of raw reads, i.e. discarding reads with bases with a Phred Score <30 (-fastq_truncqual 30) and a length <200 with usearch (Edgar 2010, v9.0.2132_i86linux32). From each sample (Mito 1-3, Complete 1-3), 100,000 reads were randomly selected with usearch and mapped against the assembled reference genomes using Bowtie (settings: Seed length 100, min insert size 200, max insert size 251, max mismatches 3, best match only). Random selection of 100,000 reads and mapping were repeated five times. After mapping, control regions were manually excluded and the number of remaining mapped sequences was determined. Enrichment factors were calculated for each species by comparing weight-corrected read numbers in the three enriched samples (Mito 1-3) to read numbers in the non-enriched samples (Complete 1-3) for all combinations (3×3). Total mitochondrial sequence enrichment was estimated by averaging enrichment factors across all combinations and species (3×3×3).

## Results

A total of 7,707,640 reads were obtained, with 1,191,468 reads for samples ‘Complete 1-3’ and 6,516,172 reads for samples ‘Mito 1-3’. After quality filtering with usearch 483,778 (Complete 1-3) respectively 2,305,181 (Mito 1-3) reads were retained. These were assembled to the complete mitochondrial genomes of *Corbicula fluminea* (17,575 bp with and 15,660 bp without control region), *Gammarus roeselii* (14,906; 13,927), *Hydropsyche exocellata* (15,789; 14,958) and *Pomphorhynchus laevis* (13,886; 13,422) (Supp1ementary material 2: Table S1, Mitogenomes: Supp1ementary material 2). Mapping of five subsets of 100,000 high quality reads each against reference genomes for which control regions were removed after mapping showed that samples not enriched for mitochondria (Complete 1-3) contained on average 0.16 % mitochondrial reads. In comparison, samples enriched for mitochondria comprised 9.47 % mitochondrial reads on average (Supp1ementary material 2: Tables S3). Species-specific enrichment factors (corrected for the relative weight per species in each sample, Supp1ementary material 2: Tables S4, S5) were 88.1 (SD: 56.6) for *Gammarus roeselii*, 163.3 (SD: 141.6) for *Hydropsyche exocellata* and 168.8 (SD: 121.4) for *Corbicula fluminea*. Overall enrichment for the whole mock communities was 140.1 (SD: 114.4) (Figure 1, Supp1ementary material 2: Table S6). Mapping against whole mitochondrial genomes including control regions showed a 129.1-fold (SD: 92.6) enrichment of mitochondrial reads (Supp1ementary material 2: Tables S7, S8, S9). Mapping of all 2,305,181 high quality reads from samples Mito 1-3 against mitogenomes without control regions showed that *G. roeselii* was sequenced with a mean coverage of 1028.8 ± 182.1, *H. exocellata* with 1109.9 ± 371.7, *C. fluminea* with 1114.4 ± 283.5 and *Pomphorhynchus laevis* with a coverage of 132.7 ± 46.1 (Coverage calculated with Geneious, Supp1ementary material 2: Table S10). Read numbers per sample are listed in Table S11 (Supp1ementary material 2) and reads have been deposited in the SRA.

**Figure 1:**
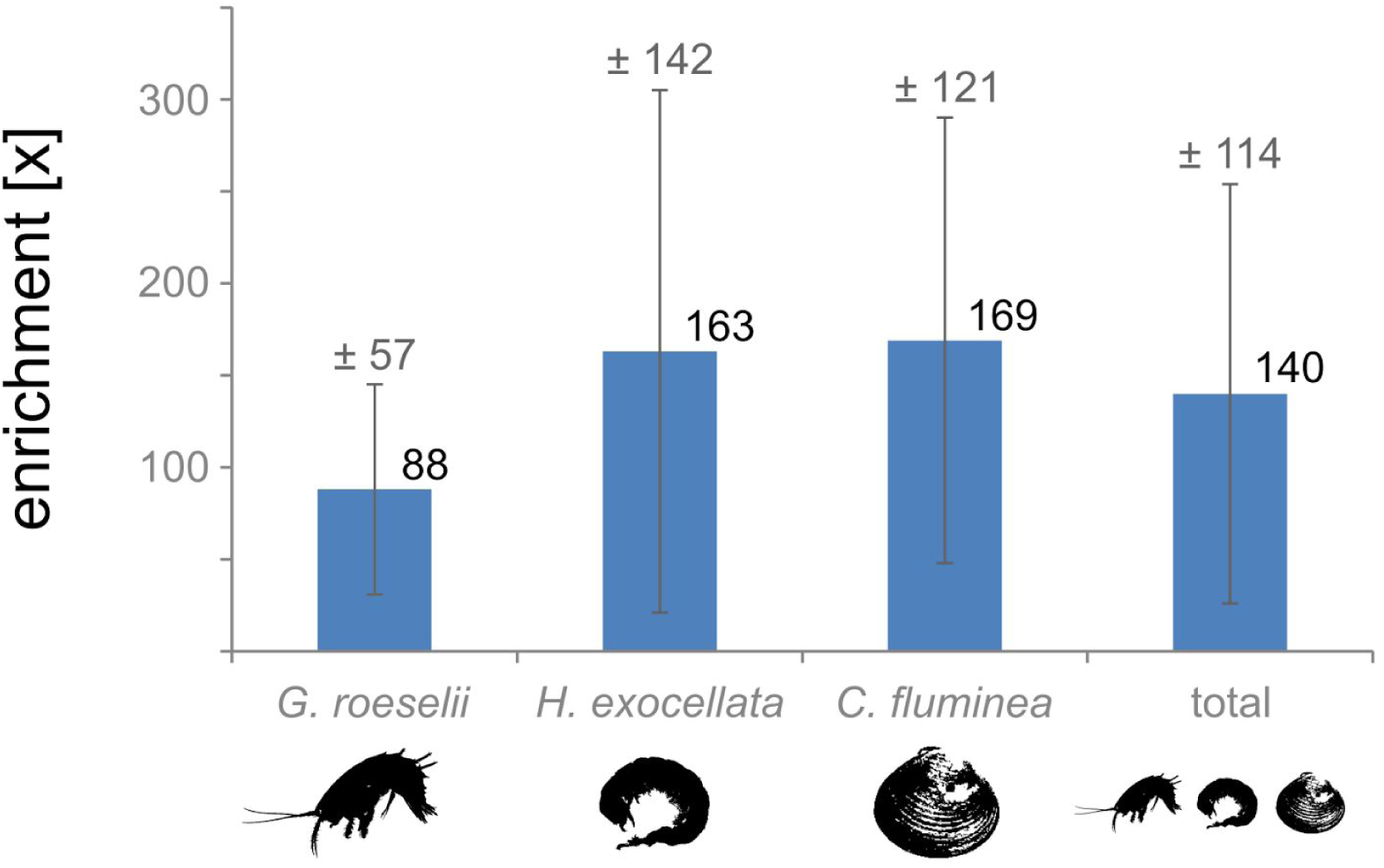
Mitochondrial DNA enrichment factor and standard deviation for the studied species and the complete mock community.

## Discussion

We developed and tested a simple centrifugation protocol for the enrichment of mitochondrial DNA from ‘mock’-communities. By using this technique and sequencing 6,516,172 raw reads for samples enriched for mitochondria, the full mitochondrial genomes of the amphipod *Gammarus roeselii*, the caddisfly *Hydropsyche exocellata* and the clam *Corbicula fluminea* were sequenced. In addition, we recovered the mitogenome of the acanthocephalan endoparasite *Pomphorhynchus laevis*, being present in *G. roeselii*.

We demonstrated that samples enriched for mitochondria contained on average 9.47 % high quality mitochondrial reads, while samples not enriched for mitochondria contained on average 0.16 %. Overall, the enrichment was 140-fold when mitogenomes without control regions were analyzed, and 129-fold when control regions were included in the analyses. However, standard deviations are high (114 with control region excluded and 93 with control region included), which is likely due to unequal homogenization of specimens and subsequent uneven release and enrichment of mitochondria and because enrichment was performed on other communities than for the controls. The weight-adjusted calculations of enrichment factors via permutations thus reflect a proxy but likely overestimates variation because sampling of control and treatment used different communities, albeit from the same population/streams. The minimum mean coverage for one of the target species was 1028.8 ± 182.4 (*Gammarus roeselii*) and still 132.7 ± 46.1 for the acanthocephalan parasite *Pomphorhynchus laevis*, when only 2,305,181 high quality reads obtained from samples enriched for mitochondria were analyzed. This highlights the great potential of the applied technique for fast sequencing of whole mitochondrial genomes, as even the relatively small parasite species was sequenced with an appropriate coverage. By using our protocol, bulk samples can be expected to be sequenced with a high enough coverage that allows the detection of mitochondrial sequences for many hundreds to thousands of specimens in parallel. However, taxonomic assignments of these reads require the completion of reliable, well-curated full mitochondrial reference libraries. Our technique for mitochondrial enrichment can greatly speed up this process, and it can be expected to obtain full mitochondrial genomes of at least several dozens of species with a single MiSeq lane if specimens are carefully selected and varying biomasses are accounted for (Elbrecht *et al*. 2017). The latter step is important as specimens with a high biomass can potentially prevent smaller specimens from being sequenced with a high enough coverage. Recovery of diverse taxonomic groups from mitogenomic datasets has been shown to work well with samples not enriched for mitochondrial DNA (Gillett *et al*. 2014; Arribas *et al*. 2016), and efficiency is expected to greatly increase by the application of mitochondrial enrichment. Although our protocol leads to a high enrichment of mitochondrial DNA, results suggest that the technique can be further improved in order to make the procedure of extracting mitochondria from tissue more reliable and standardised. We propose an automated homogenisation technique using machines instead of manual homogenisation with mortar and pestle, which is thought to often not crack cells and release intact mitochondria. A refined technique is expected to lead to an even higher and more even enrichment of mitochondrial DNA – a desirable goal since still around 90 % of all produced reads are of putative nuclear origin and enrichment is uneven for different species. Also, further studies addressing the enrichment efficiency for different species and the biomass to reads ratio are needed to further explore the potential of mitogenomic approaches. Despite this, our study demonstrates that the application of a simple centrifugation protocol enriches mitochondrial DNA 140-fold on average from mock-communities containing several species. The achieved coverage of complete mitochondrial genomes of minimum 1028.8 for our target species from 2.3 million sequences makes it obvious that even with 10 % of resulting high quality mitochondrial reads, many specimens could be sequenced and their mitogenomes assembled in a single MiSeq run. The unintentional discovery and effective mitogenome assembly of the acanthocephalan parasite *Pomphorhynchus laevis* further strengthens the approach, as PCR primers and probes often do not capture unexpected taxa for which primers or probes have not been designed. Finally, our protocol can easily be applied in the field if a cooling centrifuge can be transported to a nearby location, allowing to process the fresh tissue material needed for high(er) rates of mitochondrial enrichments. The ease of application in combination with a) a minimized laboratory workload, b) greatly reduced costs compared to mitogenomic approaches without mitochondrial enrichment and c) the high sequencing coverage per recovered mitogenome renders our mitochondrial enrichment protocol ideal for the fast generation of reference libraries (‘mitogenome skimming’) and subsequently also for biodiversity assessments.

## Acknowledgments

We thank Cristina Hartmann-Fatu for help in the lab and the Herzog-Sellenberg Foundation for financial support. This article is based upon work from COST Action DNAqua-Net (CA15219), supported by the COST (European Cooperation in Science and Technology) program.

## Author contributions

JNM, VZ, AMW and FL designed the study. JNM and VZ sampled the specimens and performed laboratory work. JNM, VZ, AMW, FL analysed the data. JNM, VZ, AMW, FL wrote the manuscript. All authors read and approved the final version of the manuscript.

